# CaCO_3_ Nanoparticles Delivering MicroRNA-200c Suppress Oral Squamous Cell Carcinoma

**DOI:** 10.1101/2023.10.05.561110

**Authors:** Qiong J Ding, Matthew T. Remy, Chawin Upara, Jue Hu, Andrés V. Mora Mata, Amanda J. Haes, Emily Lanzel, Hongli Sun, Marisa R. Buchakjian, Liu Hong

## Abstract

MicroRNA (miR)-200c suppresses the initiation and progression of oral squamous cell carcinoma (OSCC), the most prevalent head and neck cancer with high recurrence, metastasis, and mortality rates. However, *miR-200c*-based gene therapy to inhibit OSCC growth and metastasis has yet to be reported. To develop an miR-based gene therapy to improve the outcomes of OSCC treatment, this study investigates the feasibility of plasmid DNA encoding *miR-200c* delivered via non-viral CaCO_3_-based nanoparticles to inhibit OSCC tumor growth. CaCO_3_-based nanoparticles with various ratios of CaCO_3_ and protamine sulfate (PS) were utilized to transfect pDNA encoding *miR-200c* into OSCC cells and the efficiency of these nanoparticles was evaluated. The proliferation, migration, and associated oncogene production, as well as *in vivo* tumor growth for OSCC cells overexpressing *miR-200c* were also quantified. It was observed that, while CaCO_3_-based nanoparticles improve transfection efficiencies of pDNA *miR-200c*, the ratio of CaCO_3_ to PS significantly influences the transfection efficiency. Overexpression of *miR-200c* significantly reduced proliferation, migration, and oncogene expression of OSCC cells, as well as the tumor size of cell line-derived xenografts (CDX) in mice. In addition, a local administration of pDNA *miR-200c* using CaCO_3_ delivery significantly enhanced *miR-200c* transfection and suppressed tumor growth of CDX in mice. These results strongly indicate that the nanocomplexes of CaCO_3_/pDNA *miR-200c* may potentially be used to reduce oral cancer recurrence and metastasis and improve clinical outcomes in OSCC treatment. (227 words)

## Introduction

Oral squamous cell carcinoma (OSCC) is a highly invasive cancer and accounts for approximately 90% of oral cavity malignancies (Bagan et al. 2010). Although postoperative adjuvant radiotherapy or chemoradiotherapy are utilized for advanced OSCC tumors, nearly 40% of patients with advanced stage cancer will experience a recurrence. The 5-year survival rate for advanced stage OSCC is less than 50% (Adel et al. 2016; Chinn and Myers 2015; Wang et al. 2013).

MicroRNAs (miRs) are small non-coding RNAs that play crucial roles in OSCC initiation and progression via epigenetically regulating cellular biological processes (Aali et al. 2020; Emfietzoglou et al. 2020; Manzano-Moreno et al. 2021; Momen-Heravi and Bala 2018; Troiano et al. 2018; Yoshizawa and Wong 2013). miR-200 family members are highly involved in OSCC (Arunkumar et al. 2018; Hsieh et al. 2021; Kim et al. 2019). Specifically, the levels of *miR-200c* in OSCC tumor tissues and the circulating exosomes are significantly less than in adjacent normal tissues (Song et al. 2020). The downregulated level of *miR-200c* in tumor tissues correlates with OSCC treatment outcomes. In addition, *miR-200c* suppresses OSCC stemness and tumor initiating properties by targeting Sox2/Wnt signaling (Liu et al. 2017), and attenuates OSCC aggressiveness-correlated inflammation and progression by targeting NF-ĸB signaling and epithelium-mesenchymal-transition (EMT) (Brabletz et al. 2011; Johnson et al. 2016; Sztukowska et al. 2016; Tamagawa et al. 2014). *miR-200c* can also inhibit the proliferation of OSCC cells (Yan et al. 2018). In addition, *miR-200c* reduces chemoradiation resistance in oral cancer treatment by targeting the hub genes involved in regulating cisplatin and docetaxel resistance in OSCC (Brabletz et al. 2011; Cui et al. 2020; Wu et al. 2019). These previously published studies strongly indicate that overexpressing *miR-200c* would improve OSCC treatment effectiveness by preventing tumor recurrence and metastasis. However, *miR-200c*-based gene therapy for OSCC treatment has not yet been studied.

CaCO_3_ is a well-studied, mineral biomaterial with excellent osteoconductivity, biocompatibility, and biodegradability in bone formation. Nanosized FDA-approved CaCO_3_/DNA co-precipitates were also developed for gene delivery (Sharma et al. 2015). Notably, protamine sulfate (PS), a clinically used antidote for heparin-induced anticoagulation, has been reported to substantially improve the transfection efficiency of CaCO_3_-mediated plasmid DNA (pDNA) into cells (He et al. 2018; Wang et al. 2014). Our previous studies have demonstrated that pDNA *miR-200c* delivered by CaCO_3_ effectively improves transfection efficiency and alveolar bone formation (Remy et al. 2022). In the present studies, the transfection efficiencies of pDNA *miR-200c* into OSCC cells were investigated by varying ratios of CaCO_3_ and PS, and anti-OSCC capacities of pDNA encoding *miR-200c* were evaluated subsequently *in vitro* and *in vivo*. It is demonstrated that CaCO_3_-based nanoparticles effectively transfect pDNA *miR-200c* into OSCC cells and significantly inhibit the viability and migration of OSCC cells and reduce tumor growth in mice. This evidence strongly indicates that the nanoparticles composed of FDA-approved CaCO_3_ and PS may deliver pDNA encoding *miR-200c* to prevent tumor recurrence and metastasis in clinical OSCC treatment.

## Materials and Methods

1. *Preparation and characterization of pDNA miR-200c and CaCO_3_ nanocomplexes with different ratios of CaCO_3_ and PS*. pDNA encoding *miR-200c*, empty vector (EV), and scramble vector (SV) were prepared as in our previous studies (Hong et al. 2016). CaCO_3_/pDNA *miR-200c* nanocomplexes at CaCO_3_:PS ratios of 1:0.03125, 1:0.125, 1:0.25, and 1:0.5 were prepared as previously described (Remy et al. 2022). The nanocomplexes were visualized using scanning electron microscopy (SEM; Hitachi S-4800, Japan). The hydrodynamic particle diameter and zeta potential of the CaCO_3_/pDNA nanocomplexes were measured using dynamic light scattering (DLS) (Zetasizer Nano ZS; Malvern Instruments, Worcestershire, UK) as previously described (Remy et al. 2022).
2. *Transfection of pDNA miR-200c into OSCC cells using CaCO_3_-based nanoparticles with different ratios of CaCO_3_:PS*. To determine the influence of CaCO_3_:PS ratio on *miR-200c* transfection efficiency, 2×10^4^ OSCC cells (SCC 193 cells; Millipore Sigma, Burlington, MA) were treated with 1 µg pDNA encoding *miR-200c*/CaCO_3_ nanocomplexes at different ratios of CaCO_3_:PS in Opti-MEM®. To determine the transfection efficiency of CaCO_3_/*miR-200c* compared to other delivery systems, we transfected SCC 193 cells with 1 μg pDNA encoding empty vector (EV) or *miR-200c* using either naked pDNA alone, branched PEI (MW 25 kDa; Sigma-Aldrich, St. Louis, MO), or CaCO_3_/PS co-precipitated nanoparticles. The PEI/pDNA *miR-200c* (3:1 ratio) was prepared according to previous studies (Hong et al. 2016). All *in vitro* transfections were up to 16 hours and then treated with fresh DMEM high glucose medium supplemented with 10% fetal bovine serum (FBS). *miR-200c* transfection efficiency was assessed via qRT-PCR.
3. *Cell viability, migration, and oncogenic marker measurements after pDNA encoding miR-200c transfection*. SCC 180 and 193 cells (Millipore Sigma, Burlington, MA) were used to investigate OSCC cell viability, migration, and oncogenic marker expression after treatment with pDNA *miR-200c*/CaCO_3_ nanocomplexes. To determine the viability of OSCC cells, 2×10^3^ cells were seeded in 96-well plates and then treated with 0.2 μg *miR-200c*/CaCO_3_. The cell viability was documented via 3-(4,5-dimethylthiazol-2-yl)-2,5-diphenyltetrazolium bromide (MTT) assay (Biotium, Fremont, CA). To investigate cell migration, 2×10^4^ cells were cultured to confluency in 12-well plates. After 2 μg pDNA *miR-200c*/CaCO_3_ nanocomplexes or the same amount of EV and SV were added, a pipette tip was used to create a linear wound scratch across the confluent cell monolayer. Images were captured daily and cell migration motility was measured using the repaired area (ImageJ). To investigate the inhibitory function of *miR-200c* transfection on OSCC cells, 10^4^ cells were treated with 1 μg plasmid *miR-200c*/CaCO_3_ nanocomplexes. Total RNA of the cells transfected with *miR-200c* and the same amount EV were extracted after three days. Oncogenic markers of OSCC, including Neurogenic locus notch homolog protein 1 (*Notch1*), FAT atypical cadherin 1 (*FAT1*), and Cyclin Dependent Kinase Inhibitor 2A (*CDKN2A*), were quantitatively measured using qRT-PCR.
4. *In vivo tumor growth of OSCC cells pretreated with pDNA miR-200c/CaCO_3_ nanocomplexes*. All proposed animal studies were performed following a protocol approved by the Institutional Animal Care and Use Committee (IACUC) at the University of Iowa. 2×10^6^ SCC 193 cells were treated with 20 μg *miR-200c*/CaCO_3_ nanocomplexes or the same amount of EV/CaCO_3_ complexes overnight. The cells suspended in 100 μl of DMEM were then subcutaneously injected into flanks of eight-week-old male BALB/c nude mice (Jackson lab). The tumors were harvested and analyzed after five weeks. Total RNA was extracted from half of the tumors to measure osteogenic markers. The other half was stained with hematoxylin & eosin (H&E) and immunofluorescence (IF) for histological analyses. The histological analysis of explant section after H&E staining was performed blindly by a pathologist.
5. *Local administration of pDNA miR-200c/CaCO_3_ nanocomplexes inhibiting OSCC tumor growth in vivo*. 2×10^6^ SCC 193 cells were suspended in 50 μl of serum-free DMEM and injected subcutaneously into the flanks of nude mice. After 24h hours, 20 μg pDNA *miR-200c*/CaCO_3_ complexes or EV/CaCO_3_ controls were suspended in 100 μl solution and injected into the site of OSCC cell injection. Mice were sacrificed after three weeks, and the tumors were analyzed using qRT-PCR and immunohistochemistry. The histological analysis of explant sections after H&E staining was performed blindly by a pathologist.
6. *qRT-PCR*. Total RNA was extracted with the Qiagen miRNeasy microKit (Qiagen, Hilden, Germany). miR extracted from the samples was reverse transcribed into cDNA using Takara kit (Otsu, Shiga, Japan). cDNA was used to amplify miRNA and the reference gene *U6* via PCR. MirX (Takara) CDNA synthesis kits were used to detect miRNA expression. The corresponding PCR primers are listed in Supplementary Table 1. Quantitative PCR was performed with the SYBR Premix Ex TaqTM II (Takara) with the CFX96 Real-Time PCR System (Bio-Rad, Hercules, CA). Comparative real-time PCR was performed in replicate and relative expression was obtained using the comparative Ct (ΔΔCt) method.
7. *Immunofluorescence (IF) and H&E staining*. Tumor tissues were fixed in 4% paraformaldehyde overnight and immersed in 18% sucrose. The tissues were embedded in optimal cutting temperature (OCT) compound and cut into 7 μm sections. For IF staining, the slides were incubated with primary antibody, including CDKN2A, FAT1, NOTCH1 (1:50 dilution), overnight at 4°C, and with fluorescently labeled secondary antibody goat anti-rabbit 488 antibody (1:100 dilution) (Elabscience, Houston, TX). Samples were then mounted in ProLong Gold Antifade Mountant with 4,6-diamidino-2-phenylindole (DAPI) (Thermo Fisher Scientific, MA). Images were acquired using a microscope (Eclipse Ts2; Nikon, Melville, NY) with constant acquisition parameters. The frozen sections (7 μm) were also stained with H&E.
8. *Statistical analysis*. Descriptive statistics were conducted for both *in vitro* and *in vivo* investigations. One-way ANOVA with post-hoc Tukey’s honestly significant difference (HSD) tests were used to determine significant differences between treatment groups for the *in vitro miR-200c* transfection and *in vivo* tumors. A two-tailed unpaired Student’s t test was used for comparing tumor growth of OSCC cells with *miR-200c* and EV transfection. All statistical tests used a significance level of 0.05, and each graphic depicts mean values and standard deviations (SD).

## Results

*CaCO_3_-based nanoparticles enhance transfection efficiencies of pDNA miR-200c in OSCC cells.* Figure 1A summarizes the expression level of *miR-200c* after pDNA *miR-200c* delivery by CaCO_3_-based nanoparticles and other delivery systems. Notably, naked pDNA *miR-200c* increases expression of *miR-200c* more substantially than the EV and untreated controls; yet CaCO_3_/pDNA *miR-200c* significantly increases *miR-200c* expression in comparison to naked pDNA transfection. Although PEI transfection also increased *miR-200c* more than naked pDNA and untreated controls, *miR-200c* expression after PEI transfection was significantly lower than pDNA *miR-200c* delivered by CaCO_3_-nanoparticles at CaCO3:PS ratio of 1:0.25. However, EV delivered by CaCO_3_ had no ability to increase *miR-200c* expression. The expression level of *miR-200c* is dose-dependent based on the doses of CaCO_3_/pDNA *miR-200c* nanocomplexes (Figure 1B), and continuously increased after six days of transfection (Figure 1C). Among the different ratios of CaCO_3_:PS, *miR-200c* expression was significantly highest when delivered by CaCO_3_/PS at 1:0.25 than at other CaCO_3_/PS ratios (Figure 1D).

**Figure 1.**
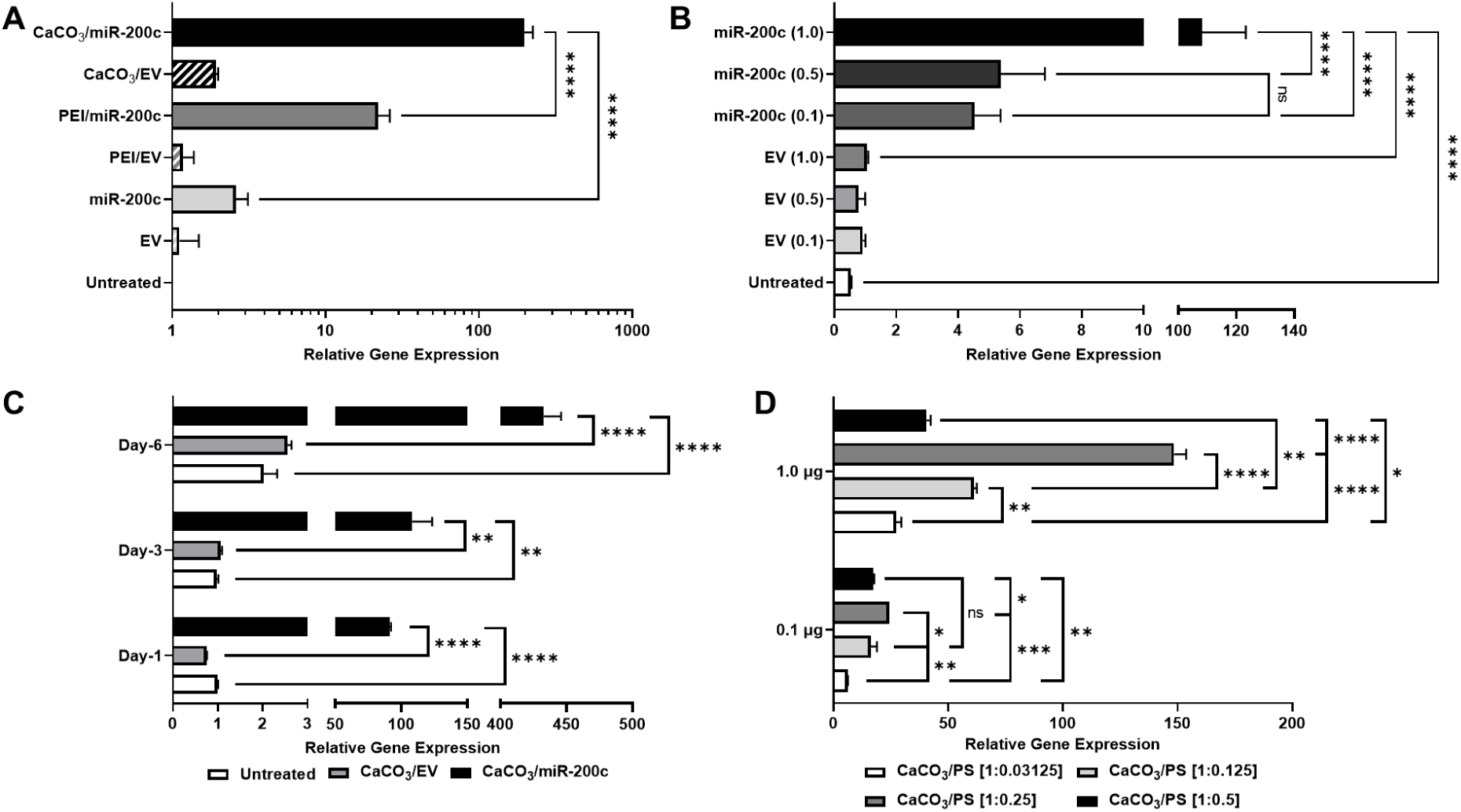
CaCO_3_-based nanoparticles improve the transfection efficiency of pDNA *miR-200c* in OSCC cells. (A) Normalized *miR-200c* transcript in SCC 193 treated with pDNA *miR-200c* and EV at 1.0 µg delivered by naked pDNA, PEI, and CaCO_3_ nanoparticles; (B) Normalized *miR-200c* transcripts in SCC 193 cells with CaCO_3_ nanoparticle delivered pDNA *miR-200c* and EV at different concentrations (from 0.1 to 1.0 μg); (C) Normalized *miR-200c* transcripts in SCC 193 cells with CaCO_3_ nanoparticle delivered 1.0 µg pDNA *miR-200c* and EV after different times; (D) Normalized transcripts of *miR-200c* delivered by CaCO_3_-based nanoparticles with different ratios of CaCO_3_:PS. *: p<0.05, ****: p<0.0001.

### CaCO_3_:PS ratios affect characteristics of CaCO_3_/pDNA miR-200c nanocomplexes

Figure 2A and B summarize the SEM images and quantitative diameter measurements of the nanocomplexes at different CaCO_3_:PS ratios. There is notably an inverse relationship between size and PS concentration, where the diameter of the nanocomplexes decreases with increasing amounts of PS. The size of CaCO_3_:PS at 1:0.25 (128 ± 32 nm) and 1:0.5 (74 ± 33 nm) are significantly smaller than the CaCO_3_:PS at 1:0.125 (433 ± 92) and 1:0.03125 (3312 ± 1055 nm) (Figure 2B). In addition, the nanoparticles at 1:0.03125 and 1:0.125 ratio of CaCO_3_:PS exhibit irregular crystalline morphologies, while the particles at a higher ratio of CaCO_3_:PS are structurally round (Figure 2A). Under hydrodynamic conditions, we also confirm that the diameters of nanocomplexes made by CaCO_3_:PS at 1:0.25 (176 ± 50 nm) and 1:0.5 (178 ± 16 nm) are significantly smaller than the CaCO_3_:PS at 1:0.125 (341 ± 0 nm) and 1:0.03125 (367 ± 56 nm) (Figure 2C). However, the zeta potential surface was not statistically different among the nanocomplexes at different CaCO_3_:PS ratios (Figure 2D).

**Figure 2.**
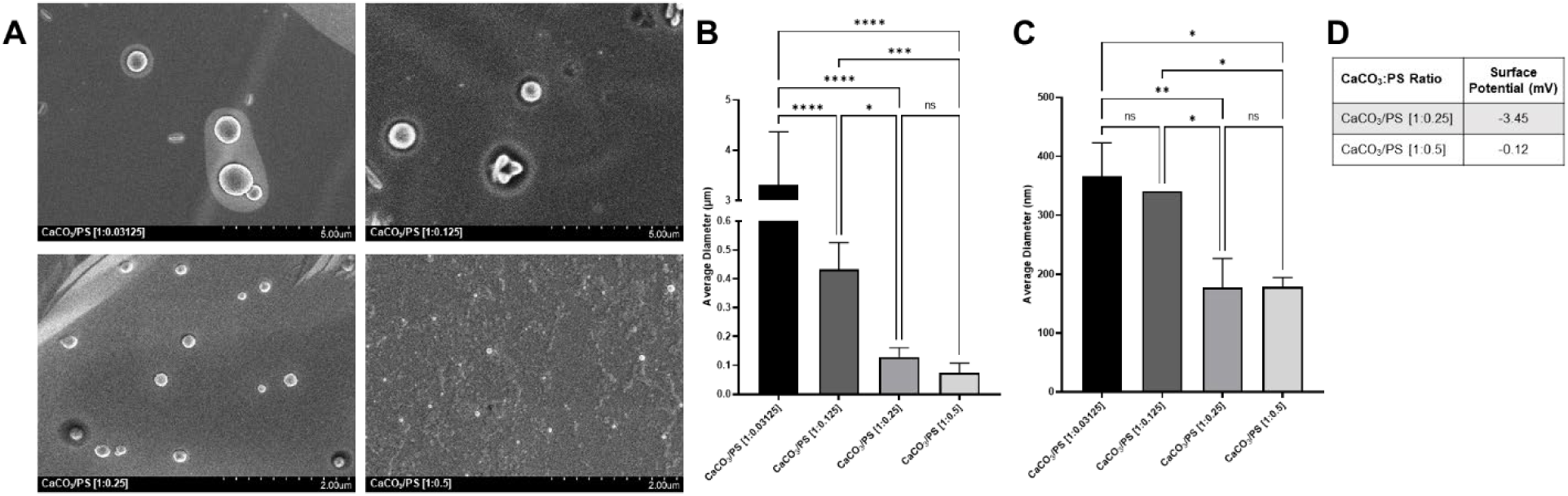
Characterization of pDNA *miR-200c*/CaCO_3_ nanocomplexes. SEM images (A), diameters measured by SEM images (B) and DLS (C), and zeta surface potential of pDNA *miR-200c*/CaCO_3_ with different ratios of CaCO_3_:PS (D). *: p<0.05, **: p<0.01, ***: p<0.001, ****: p<0.0001.

### Transfection of pDNA encoding miR-200c using CaCO_3_-based nanoparticles reduces viability, migration, and oncogene expression of OSCC cells in vitro

Two OSCC cell types were treated with CaCO_3_/pDNA encoding *miR-200c* at a CaCO_3_:PS ratio of 1:0.25. The viabilities of OSCC cells measured by MTT assay after treatment with pDNA encoding *miR-200c* are significantly lower than that of untreated OSCC cells. The same dose of EV and SV has no inhibitory effect on the viability of OSCC cells (Figure 3A, B). In addition, the treatment of CaCO_3_/*miR-200c* significantly inhibited cancer cell migration using a scratch assay. The non-cell scratch areas remained open after one and two days for SCC 192 and SCC 180 cells, respectively, while untreated cells and cells treated with EV and SV quickly migrated and covered the scratched area (Figure 3C, D). Oncogenic markers associated with OSCC cells, including *Notch1*, *FAT1*, and *CDKN2A*, were significantly downregulated in cells treated with CaCO_3_/*miR-200c* (Figure 3E). In addition, OSCC oncogenes, including PIK3CA (Phosphatidylinositol-4,5-Bisphosphate 3-Kinase Catalytic Subunit Alpha), Tumor protein P53, Neurogenic locus notch homolog protein 1 (Notch1), FADD (Fas Associated Via Death Domain) and CCND (Cyclin D1), were downregulated by miR-200c overexpression (see Supplement Files).

**Figure 3.**
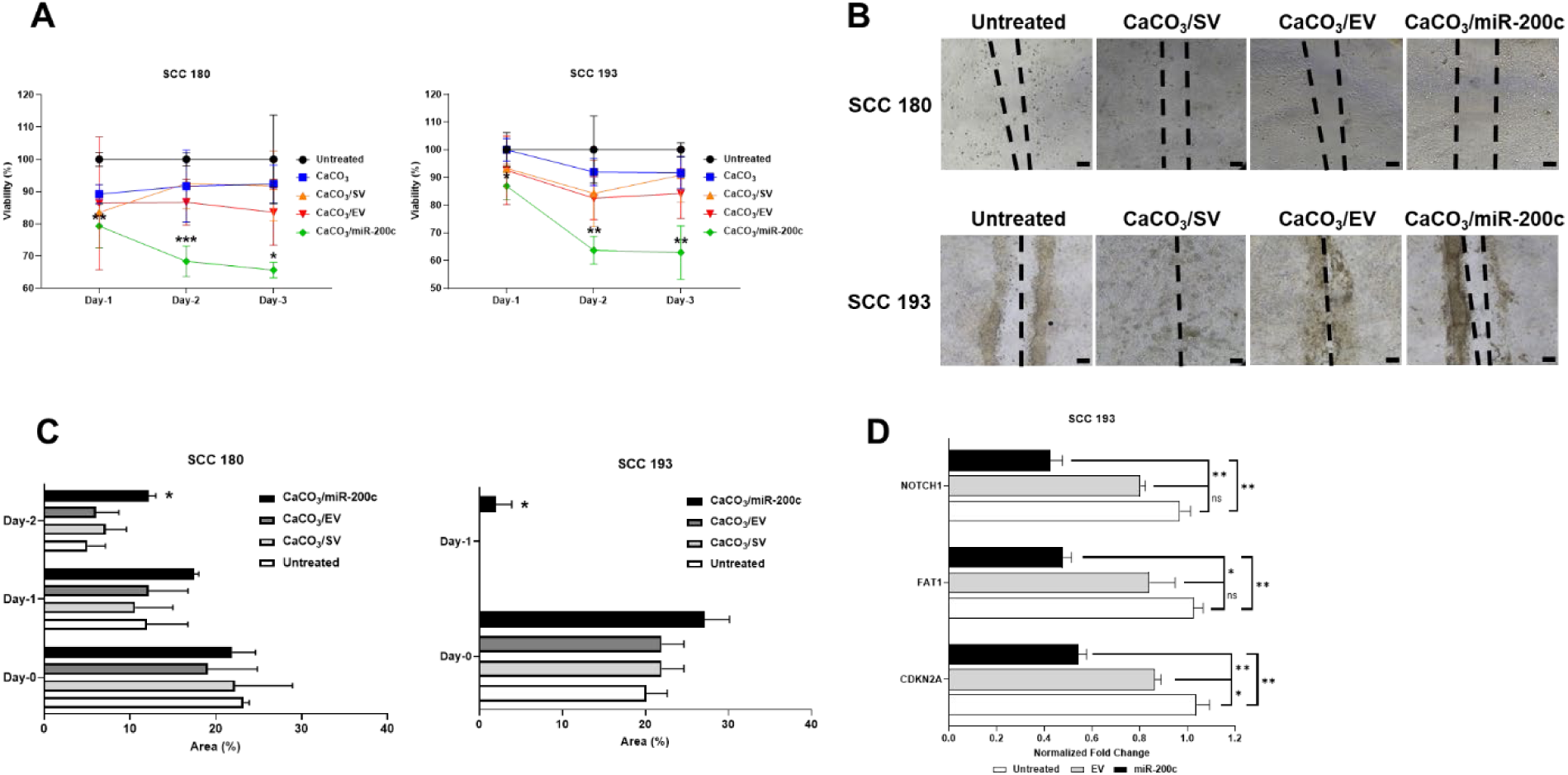
Oncogenic characterization of OSCC cells treated with pDNA *miR-200c*/CaCO_3_ nanocomplexes and controls. (A) Viabilities of SCC 180 and SCC 193 cell transfected with 0.1 µg pDNA *miR-200c*, EV, and SV delivered by CaCO_3_-based nanoparticles; (B) Photographs 24 hrs after a wound-healing/scratch assay of OSCC cells transfected with 0.1 µg pDNA *miR-200c* and controls; (C) Quantitative measurement of un-cell area percentages after a wound-healing/scratch assay of OSCC cells with different treatment; (D) Normalized fold changes of OSCC associated oncogene transcripts of SCC 193 transfected with pDNA *miR-200c* and controls after 3 days. *: p<0.05, **: p<0.01, ***: p<0.001 vs untreated and EV/SV. Scale bars: 100 µm.

### Overexpression of miR-200c with CaCO_3_/miR-200c nanocomplex treatment reduces OSCC cell-derived tumor growth in vivo

To understand the tumor growth inhibition of OSCC cells after transfection with pDNA encoding *miR-200c*, CaCO_3_/pDNA encoding *miR-200c* were transfected to SCC 193 cells overnight and then injected subcutaneously to the flank of BALB/c nude mice. After five weeks, the harvested tumor weight induced by SCC 193 pretreated with CaCO_3_/pDNA *miR-200c* was significantly smaller than those with EV and SV (Figure 4A, B). The transcripts of OSCC associated oncogenes, including *CDKN3A*, *FAT1*, and *NOTCH1*, were significantly downregulated in tumors induced by cells with *miR-200c* overexpression, compared to the cells pretreated with EV (Figure 4C). Immunofluorescent stains using antibodies against CDKN2A, FAT1, and NOTCH1 showed the positive stains in sections of tumor cells pretreated with EV and SV. However, limited positive staining was found in the tumors of cells pretreated with CaCO_3_/pDNA *miR-200c* nanocomplexes (Figure 4D). In addition, according to histological examination by pathologists, the morphology of cells in explants of OSCC cells pretreated with *miR-200c* show no evidence of SCC cells, while the cells in OSCC cells treated with EV or SV are well-differentiated or moderately-differentiated SCC cells.

**Figure 4.**
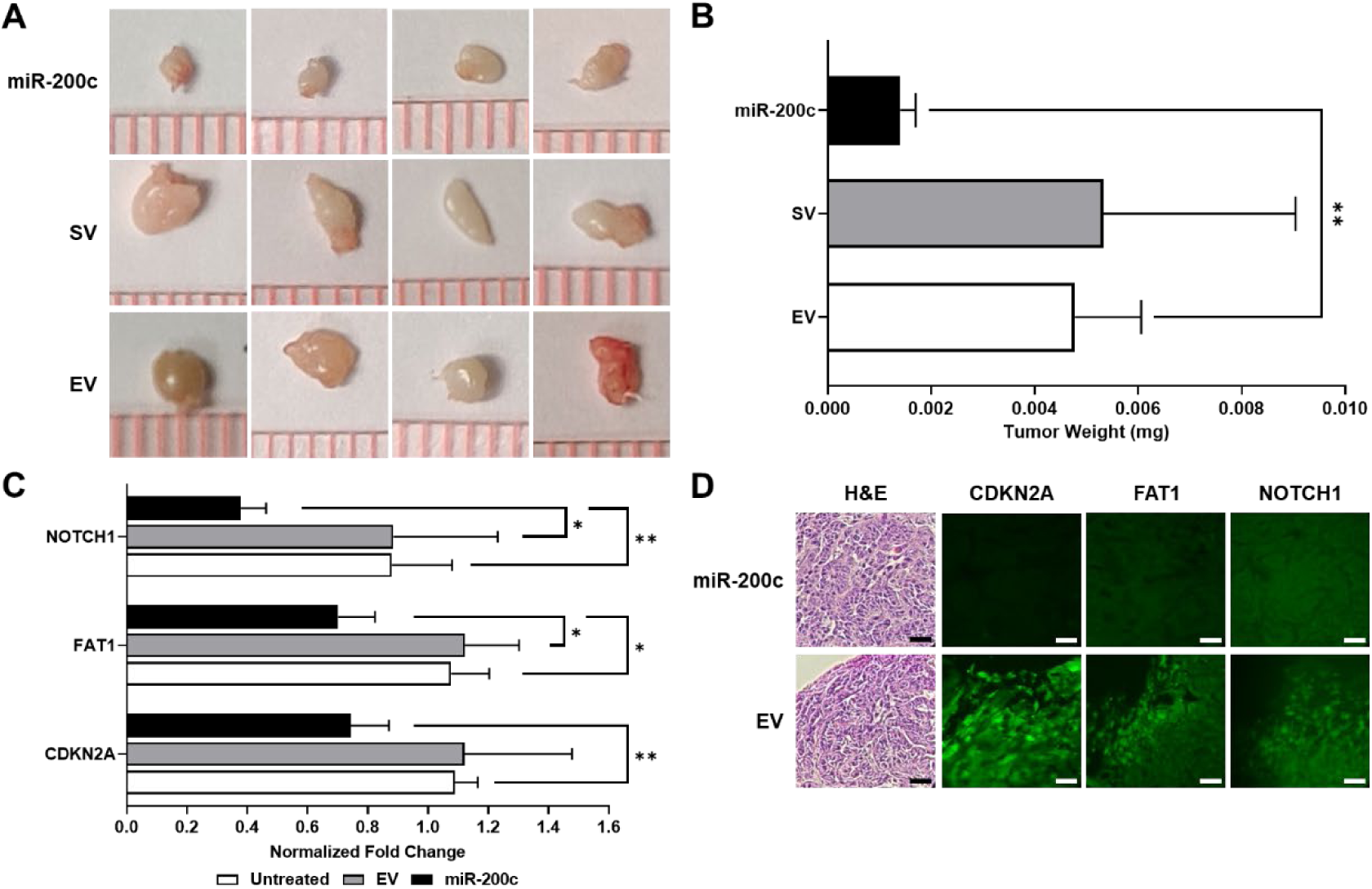
Transfection of pDNA *miR-200c* by CaCO_3_-based nanoparticles suppresses tumor growth of OSCC *in vivo*. (A,B): Photographs of OSCC tumor explants (A) and the weight of tumors (B) induced by SCC 193 cells pretreated with CaCO_3_/*miR-200c* complexes, CaCO_3_/EV, or CaCO_3_/SV and subcutaneously implanted into the flank of nude mice for 5 weeks. (C) Normalized transcript fold changes of OSCC-related oncogenes of tumor explants with different treatments. (D) Microphotographs of sections of OSCC tumor explants induced with the OSCC cells with different treatments after stained with H&E and immunofluorescence with antibodies against CDKN2A, FAT1, and NOTCH. All biostatistical analysis was performed by ANOVA with post-hoc Tukey’s HSD test or student’s t-test. *: p<0.05, **: p<0.01 vs EV and SV. N=4. Scale bars: 50 µm.

### Local application of CaCO_3_/pDNA miR-200c complexes inhibits OSCC tumor growth in CDX model

After a cell-derived xenograft of OSCC using SCC 193 cells was created, CaCO_3_/*miR-200c* nanocomplexes were injected to test whether local application of pDNA *miR-200c* delivered by CaCO_3_-based nanoparticles inhibit OSCC growth *in vivo*. pDNA containing GFP was used to track the transfection following the local injection. After three days, positive GFP expression in the harvested tumor mass was observed using fluorescent microscopy (Figure 5A).

**Figure 5.**
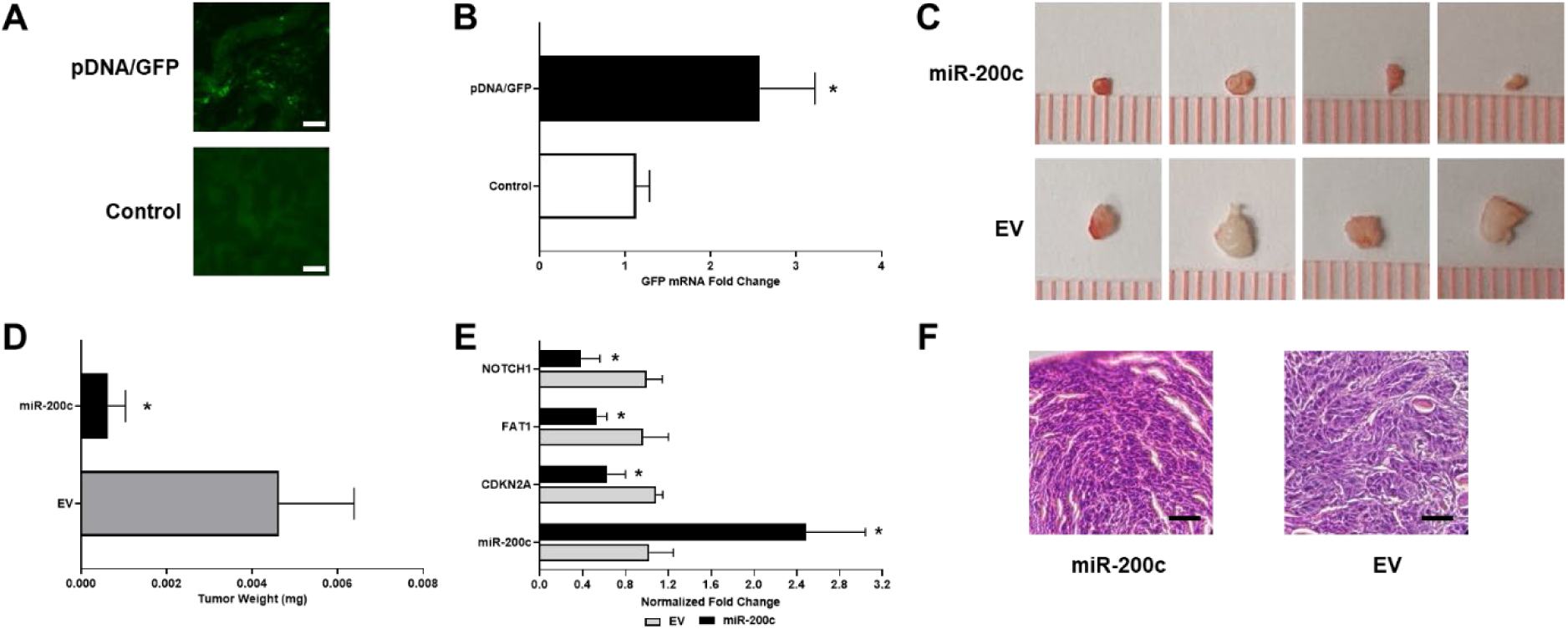
Local application of CaCO_3_/pDNA *miR-200c* effectively improves the transfection efficiency of *miR-200c* and inhibits OSCC tumor growth in a CDX model. (A,B): Immunofluorescence images and quantitative measurements of GFP expression in OSCC tumors treated with local injection CaCO_3_/pDNA encoding GFP and untreated control; (C-E): Photographs of OSCC tumor explants (C), the weight of tumors (D), and transcript fold changes of *miR-200c* and OSCC-related oncogenes within tumors (E) 3 weeks after local administration of 20 μg CaCO_3_/*miR-200c* or CaCO_3_/EV. (F) Microphotographs of sections of OSCC tumor explants induced with the OSCC cells with different treatments, H&E. *: p<0.05 vs EV/Control. N=4. Scale bars: 50 µm.

Quantitatively, GFP expression measured qRT-PCR was significantly higher in tumor tissue with *miR-200c* treatment than that of untreated controls (Figure 5B). Figures 5C and D summarize the tumor growth size after different treatments. Notably, after three weeks, the weight of harvested tumors treated with local injection of CaCO_3_/*miR-200c* nanoparticles were significantly smaller than those treated with CaCO_3_/EV at the same dose. Additionally, significantly upregulated *miR-200c* expression and downregulated transcripts of oncogenic markers in tumors treated with CaCO_3_/*miR-200c* were confirmed using qRT-PCR. Well-differentiated SCC were detected both in the explants with different treatments.

## Discussion

The present study revealed that pDNA encoding *miR-200c* delivered via biocompatible and biodegradable CaCO_3_-based nanoparticles significantly improves expression of *miR-200c* in OSCC cells, which effectively reduces the OSCC tumor growth by suppressing cell viability, migration, and oncogene expression. Moreover, local administration of CaCO_3_/pDNA *miR-200c* nanocomplexes effectively transfects pDNA into tumor cells *in vivo* and suppress the growth of OSCC cell-derived tumors. This evidence suggests that CaCO_3_/pDNA *miR-200c* nanocomplexes may potentially serve as a therapeutic tool to suppress OSCC tumor growth and recurrence clinically.

Previous studies have reported that CaCO_3_-based nanoparticles can effectively improve transfection efficiencies of pDNA encoding *miR-200c* into human preosteoblasts (Remy et al. 2022). Local application of collagen sponges incorporating CaCO_3_/pDNA *miR-200c* complexes effectively enhances bone formation in alveolar bone defects (Remy et al. 2022). In the present studies, it is confirmed that the system of CaCO_3_ with PS co-precipitation nanoparticles effectively improves transfection efficiencies of pDNA encoding *miR-200c* into OSCC cells.

Local administration of pDNA using CaCO_3_-based nanoparticle delivery also enhances the pDNA transfection and increases the expression of *miR-200c* in tumor tissues in a mouse model. Because of its biocompatibility, biodegradability, and FDA-approved materials, CaCO_3_ with co-precipitation of PS may potentially be developed for clinical application as a non-viral nanoparticle delivery system. In this study, similar to previous studies, it is observed that the transfection efficiency of pDNA encoding *miR-200c* varies in the ratio of CaCO_3_:PS in OSCC cells. Among the four different ratios, the ratio of CaCO_3_:PS at 1:0.25 is optimal to enhance the transfection efficiency for pDNA encoding *miR-200c* to the highest degree. Interestingly, our findings in the current study were partially consistent with a previous report claiming that a higher concentration of PS is associated with a higher transfection efficiency (Wang et al. 2014). We identified that the lower PS concentrations (CaCO_3_:PS ratio at 1:0.03125 and 1:0.125) have relatively lower transfection efficiencies compared to a higher PS concentration. This may be due to the relatively large size of the nanocomplexes, where increasing nanoparticle size decreases transfection efficiency(Ota et al. 2013). In this study, the diameter of nanocomplexes at lower PS concentrations are ∼400 and 3000 nm, which is significantly larger than the ideal size of nanoparticles (∼100 nm) for enhancing transfection efficiency. We found that a higher concentration of PS (CaCO_3_:PS ratio 1:0.25) can effectively reduce the size of nanoparticles and reach the ideal size to enhance transfection efficiency. However, we observed that the small diameter nanoparticles at the highest PS concentration did not enhance transfection efficiency, therefore indicating that there is an optimal PS ratio that reduces nanoparticle size while enhancing transfection efficiency. While the underlying mechanism(s) remain unknown, overdoses of PS in the CaCO_3_ nanoparticles changes the electrical environment of the nanoparticle system, likely causing decreased pDNA encapsulation efficiency. Taken together, we conclude that the ratios of CaCO_3_:PS in nanocomplexes directly affect nanoparticle size, but not surface charge, for the CaCO_3_/pDNA *miR-200c* complexes, and future *in vivo* experiments to optimize the ratio of CaCO_3_/pDNA *miR-200c* for enhanced *in vivo* transfection are needed.

In the current study, we have demonstrated that local application of pDNA *miR-200c* delivered using CaCO_3_-based nanoparticles can effectively transfect pDNA and upregulate *miR-200c* expression locally. This approach reduced the growth and oncogenic marker expression of OSCC cell-derived tumors. This evidence strongly supports the idea that overexpressing *miR-200c* by local administration at the time of reconstructive surgery will improve OSCC treatment effectiveness and prevent tumor recurrence and metastasis. Although application of chemically synthesized miR mimics may simulate the inhibitory function of *miR-200c* on OSCC, it remains unclear whether transfected *miR-200c* mimics behave similarly to endogenous *miR-200c*. Studies have shown that transient transfection of miR mimics may led to the accumulation of high molecular weight RNA species (Jin et al. 2015). To achieve mature miR functional levels, miR mimics require a high concentration, which easily induces non-specific alterations in gene expression as a side effect. In addition, sustaining a high concentration with effective overexpression level using miR mimics is challenging due to its relative instability. Therefore, the potential non-specific changes in gene expression induced by the supraphysiological levels of mature miRs and the artifactual RNA species limit the application of miRs. However, transfection of pDNA encoding miRs can simulate the biogenesis of endogenous miRs.

Transfection of pDNA encoding miRs does not lead to high molecular weight RNA species. Although the level of mature miR increased by pDNA are relatively less than the miR mimics, many studies demonstrated that they sufficiently suppress the target genes. In addition, our previous studies have demonstrated that cells with overexpression of *miR-200c* after transfection of pDNA encoding *miR-200c* may upregulate *miR-200c* levels in surrounding cells by secreting *miR-200c* enriched exosomes (Krongbaramee et al. 2021). Therefore, transfection of pDNA *miR-200c* may effectively sustain the overexpression period and enhance the inhibitory function on OSCC.

## Conclusion

We have demonstrated that pDNA encoding *miR-200c* using non-viral CaCO_3_-based nanoparticle delivery system effectively increases *miR-200c* expression in OSCC cells and suppresses oncogenic activities, thereby reducing tumor growth in a preclinical animal model. In addition to its capacities to promote bone regeneration and inhibit inflammation-associated bone resorption, *miR-200c* may serve as an adjuvant tool in reconstructive surgeries of patients with advanced OSCC to enhance bone healing and decrease tumor recurrence and metastasis. Future studies to further optimize *in vivo* transfection efficiency of *miR-200c* and its functions in preclinical animal models of OSCC reconstructive surgeries are needed.

## Acknowledgments

This research was supported by the public-private partnership (P3) of the University of Iowa Strategic Initiatives Fund and the National Institute of Dental and Craniofacial Research (NIDCR) (Grant No. R01DE026433 (LH), R01DE029159 (HS)) of the National Institutes of Health (NIH). This research was further facilitated by the IR/D (Individual Research and Development) program associated with Amanda J. Haes’s appointment at the National Science Foundation (NSF). Matthew T. Remy and Qiong J. Ding would additionally like to acknowledge the support received from the NIH/NIDCR (Grant No. F31DE031153 (MTR) and T90DE023520 (MTR, QJD)).

## Author Contributions

QJD, HS, MB, and LH designed the study. QD and JH were responsible for preparation of the CaCO3 nanocomplexes under the supervision of HS. JH, MTR, AVMM, and AJH characterized the nanocomplexes. QJD was responsible for culturing cells necessary for the in vitro transfection, release, and biocompatibility studies and subsequent qRT-PCR analyses. QID, CW, and LH conducted animal surgeries. QID and CW were responsible for all animal care, euthanasia, quantitative analyses, and histological sectioning, staining, and imaging completed for the in vivo animal studies. EL was responsible to pathological diagnosis of tumor explants. SE was responsible for pDNA preparation and characterization under the supervision of BAA. MTR conducted statistical analyses and interpretation for all collected data with technical assistance from LH. QJD, CW, MTR and LH wrote and revised the manuscript. All authors discussed the results and approved the final version of the manuscript.

## References

1. Aali M, Mesgarzadeh AH, Najjary S, Abdolahi HM, Kojabad AB, Baradaran B. 2020. Evaluating the role of micrornas alterations in oral squamous cell carcinoma. Gene. 757:144936.

2. Adel M, Liao CT, Lee LY, Hsueh C, Lin CY, Fan KH, Wang HM, Ng SH, Lin CH, Tsao CK et al. 2016. Incidence and outcomes of patients with oral cavity squamous cell carcinoma and fourth primary tumors: A long-term follow-up study in a betel quid chewing endemic area. Medicine (Baltimore). 95(12):e2950.

3. Arunkumar G, Deva Magendhra Rao AK, Manikandan M, Prasanna Srinivasa Rao H, Subbiah S, Ilangovan R, Murugan AK, Munirajan AK. 2018. Dysregulation of mir-200 family micrornas and epithelial-mesenchymal transition markers in oral squamous cell carcinoma. Oncol Lett. 15(1):649-657.

4. Bagan J, Sarrion G, Jimenez Y. 2010. Oral cancer: Clinical features. Oral Oncol. 46(6):414–417.

5. Brabletz S, Bajdak K, Meidhof S, Burk U, Niedermann G, Firat E, Wellner U, Dimmler A, Faller G, Schubert J et al. 2011. The zeb1/mir-200 feedback loop controls notch signalling in cancer cells. Embo j. 30(4):770–782.

6. Chinn SB, Myers JN. 2015. Oral cavity carcinoma: Current management, controversies, and future directions. J Clin Oncol. 33(29):3269–3276.

7. Cui J, Wang H, Zhang X, Sun X, Zhang J, Ma J. 2020. Exosomal mir-200c suppresses chemoresistance of docetaxel in tongue squamous cell carcinoma by suppressing tubb3 and ppp2r1b. Aging (Albany NY). 12(8):6756–6773.

8. Emfietzoglou R, Pachymanolis E, Piperi C. 2020. Impact of epigenetic alterations in the development of oral diseases. Curr Med Chem.

9. He XY, Liu BY, Xu C, Zhuo RX, Cheng SX. 2018. A multi-functional macrophage and tumor targeting gene delivery system for the regulation of macrophage polarity and reversal of cancer immunoresistance. Nanoscale. 10(33):15578–15587.

10. Hong L, Sharp T, Khorsand B, Fischer C, Eliason S, Salem A, Akkouch A, Brogden K, Amendt BA. 2016. Microrna-200c represses il-6, il-8, and ccl-5 expression and enhances osteogenic differentiation. Plos One. 11(8).

11. Hsieh PL, Huang CC, Yu CC. 2021. Emerging role of microrna-200 family in dentistry. Noncoding RNA. 7(2).

12. Jin HY, Gonzalez-Martin A, Miletic AV, Lai M, Knight S, Sabouri-Ghomi M, Head SR, Macauley MS, Rickert RC, Xiao C. 2015. Transfection of microrna mimics should be used with caution. Front Genet. 6:340.

13. Johnson JJ, Miller DL, Jiang R, Liu Y, Shi Z, Tarwater L, Williams R, Balsara R, Sauter ER, Stack MS. 2016. Protease-activated receptor-2 (par-2)-mediated nf-kappab activation suppresses inflammation-associated tumor suppressor micrornas in oral squamous cell carcinoma. J Biol Chem. 291(13):6936–6945.

14. Kim EJ, Kim JS, Lee S, Lee H, Yoon JS, Hong JH, Chun SH, Sun S, Won HS, Hong SA et al. 2019. Qki, a mir-200 target gene, suppresses epithelial-to-mesenchymal transition and tumor growth. Int J Cancer. 145(6):1585–1595.

15. Krongbaramee T, Zhu M, Qian Q, Zhang Z, Eliason S, Shu Y, Qian F, Akkouch A, Su D, Amendt BA et al. 2021. Plasmid encoding microrna-200c ameliorates periodontitis and systemic inflammation in obese mice. Mol Ther Nucleic Acids. 23:1204–1216.

16. Liu CM, Peng CY, Liao YW, Lu MY, Tsai ML, Yeh JC, Yu CH, Yu CC. 2017. Sulforaphane targets cancer stemness and tumor initiating properties in oral squamous cell carcinomas via mir-200c induction. J Formos Med Assoc. 116(1):41–48.

17. Manzano-Moreno FJ, Costela-Ruiz VJ, García-Recio E, Olmedo-Gaya MV, Ruiz C, Reyes-Botella C. 2021. Role of salivary microrna and cytokines in the diagnosis and prognosis of oral squamous cell carcinoma. Int J Mol Sci. 22(22).

18. Momen-Heravi F, Bala S. 2018. Emerging role of non-coding rna in oral cancer. Cell Signal. 42:134–143.

19. Ota S, Takahashi Y, Tomitaka A, Yamada T, Kami D, Watanabe M, Takemura Y. 2013. Transfection efficiency influenced by aggregation of DNA/polyethylenimine max/magnetic nanoparticle complexes. Journal of Nanoparticle Research. 15(5):1653.

20. Remy MT, Ding Q, Krongbaramee T, Hu J, Mora Mata AV, Haes AJ, Amendt BA, Sun H, Buchakjian MR, Hong L. 2022. Plasmid encoding mirna-200c delivered by caco(3)-based nanoparticles enhances rat alveolar bone formation. Nanomedicine (Lond).

21. Sharma S, Verma A, Teja BV, Pandey G, Mittapelly N, Trivedi R, Mishra PR. 2015. An insight into functionalized calcium based inorganic nanomaterials in biomedicine: Trends and transitions. Colloids Surf B Biointerfaces. 133:120–139.

22. Song J, Zhang N, Cao L, Xiao D, Ye X, Luo E, Zhang Z. 2020. Down-regulation of mir-200c associates with poor prognosis of oral squamous cell carcinoma. Int J Clin Oncol. 25(6):1072–1078.

23. Sztukowska MN, Ojo A, Ahmed S, Carenbauer AL, Wang Q, Shumway B, Jenkinson HF, Wang H, Darling DS, Lamont RJ. 2016. Porphyromonas gingivalis initiates a mesenchymal-like transition through zeb1 in gingival epithelial cells. Cell Microbiol. 18(6):844–858.

24. Tamagawa S, Beder LB, Hotomi M, Gunduz M, Yata K, Grenman R, Yamanaka N. 2014. Role of mir-200c/mir-141 in the regulation of epithelial-mesenchymal transition and migration in head and neck squamous cell carcinoma. Int J Mol Med. 33(4):879–886.

25. Troiano G, Mastrangelo F, Caponio VCA, Laino L, Cirillo N, Lo Muzio L. 2018. Predictive prognostic value of tissue-based microrna expression in oral squamous cell carcinoma: A systematic review and meta-analysis. J Dent Res. 97(7):759–766.

26. Wang B, Zhang S, Yue K, Wang XD. 2013. The recurrence and survival of oral squamous cell carcinoma: A report of 275 cases. Chin J Cancer. 32(11):614–618.

27. Wang CQ, Wu JL, Zhuo RX, Cheng SX. 2014. Protamine sulfate-calcium carbonate-plasmid DNA ternary nanoparticles for efficient gene delivery. Mol Biosyst. 10(3):672–678.

28. Wu HT, Chen WT, Li GW, Shen JX, Ye QQ, Zhang ML, Chen WJ, Liu J. 2019. Analysis of the differentially expressed genes induced by cisplatin resistance in oral squamous cell carcinomas and their interaction. Front Genet. 10:1328.

29. Yan Y, Yan F, Huang Q. 2018. Mir-200c inhibited the proliferation of oral squamous cell carcinoma cells by targeting akt pathway and its downstream glut1. Arch Oral Biol. 96:52–57.

30. Yoshizawa JM, Wong DT. 2013. Salivary micrornas and oral cancer detection. Methods Mol Biol. 936:313–324.

