## supplementary files for "CaCO_3_ Nanoparticles Delivering MicroRNA-200c Suppress Oral Squamous Cell Carcinoma"

**Table 1. Primer sequences for qRT-PCR analyses.**

| Gene | Forward Primers (5' → 3') | Reverse Primers (5' → 3') |
|---|---|---|
| *CDKN2A* | CTCGTGCTGATGCTACTGAGGA | GGTCGGCGCAGTTGGGCTCC |
| FADD | GTGGCTGACCTGGTACAAGAG | GGTAGATGCGTCTGAGTTCCAT |
| *FAT1* | CATCCTGTCAAGATGGGTGTTT | TCCGAGAATGTACTCTTCAGCTT |
| *NOTCH1* | TGGACCAGATTGGGGAGTTC | GCACACTCGTCTGTGTTGAC |
| *p53* | CCTCAGCATCTTATCCGAGTGG | TGGATGGTGGTACAGTCAGAGC |
| *PIK3CA* | GAA GCA CCT GAA TAG GCA AGT CG | GAG CAT CCA TGA AAT CTG GTC GC |

**Supplementary Figures**

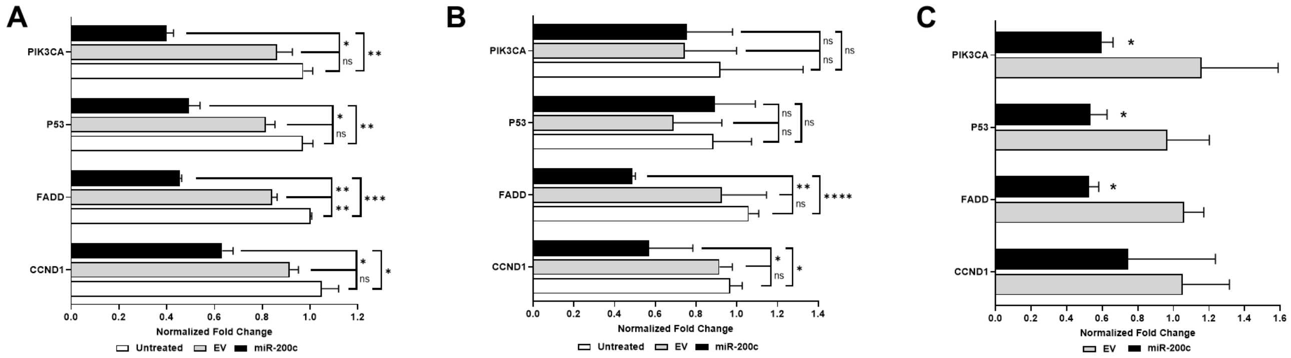


**Figure1: Oncogenic characterization of OSCC cells treated with pDNA miR-200c/CaCO3 nanocomplexes and controls *in vitro* and *in vivo*.** Normalized fold changes of *PIK3CA, P53, FADD,* and *CCND1* transcripts in SCC193 treated with pDNA encoding miR-200c *in vitro* (A), CDX of SCC 193 pretreated with pDNA *miR-200c* (B), CDX of SCC193 after treated with pDNA miR-200c *in vivo* (C).   *: $p<0.05$, **: $p<0.01$, ***: $p<0.001$. ****: $p<0.0001$.
